# QPromoters: Sequence based prediction of promoter strength in *Saccharomyces cerevisiae*

**DOI:** 10.1101/2021.04.27.441621

**Authors:** Devang Haresh Liya, Mirudula Elanchezhian, Mukulika Pahari, Nithishwer Mouroug Anand, Shivani Suresh, Nivedha Balaji, Ashwin Kumar Jainarayanan

**Affiliations:** Department of Physical Sciences, Indian Institute of Science Education and Research, Mohali, India; Department of Biological Sciences, Indian Institute of Science Education and Research, Mohali, India; Department of Computer Engineering, Ramrao Adik Institute of Technology, DY Patil Deemed to be University, Navi Mumbai, India; Department of Pharmacology, University of Oxford, Oxford OX1 3QT, UK; Kennedy Institute of Rheumatology, University of Oxford, Oxford OX3 7FY, UK; Interdisciplinary Bioscience Doctoral Training Program and Exeter College, University of Oxford, Oxford OX3 7DQ, UK

**Keywords:** *Saccharomyces cerevisiae*, promoter strength, minimal genome, linear regression, QPromoters

## Abstract

Promoters play a key role in influencing transcriptional regulation for fine-tuning expression of genes. Heterologous promoter engineering has been a widely used concept to control the level of transcription in all model organisms. The strength of a promoter is mainly determined by its nucleotide composition. Many promoter libraries have been curated but few have attempted to develop theoretical methods to predict the strength of promoters from its nucleotide sequence.

Such theoretical methods are not only valuable in the design of promoters with specified strength, but are also meaningful to understand the mechanism of promoters in gene transcription. In this study, we present a theoretical model to describe the relationship between promoter strength and nucleotide sequence in *Saccharomyces cerevisiae*. We infer from our analysis that the −49 to 10 sequence with respect to the Transcription Start Site represents the minimal region that can be used to predict the promoter strength. We present an online tool https://qpromoters.com/ that takes advantage of this fact to quickly quantify the strength of the promoters.

## Introduction

*Saccharomyces cerevisiae (S. cerevisiae)*, commonly known as brewer’s yeast, is a widely used eukaryotic model organism in synthetic biology – it has applications in the production of biofuels, recombinant proteins and bulk chemicals (1,2). Promoters are basic transcriptional elements that play a key role in manipulating genetic and metabolic pathways in *S. cerevisiae* by the regulation of protein expression both quantitatively and temporally (3,4). Promoters in *S. Cerevisiae* have multiple components which together account for successful transcriptional regulation. The key components of a yeast promoter are an upstream activator sequence (UAS), an upstream repressor sequence (URS), a nucleosome-disfavoring sequence and a core promoter region.

The core promoter is the DNA sequence nearest to the start codon, which interacts with RNA polymerase II (pol-II) and other general transcriptional factors to form the pre-initiation complex (PIC) (2). The core region also contains the TATA box, the transcription start site (TSS), a PIC localization stretch and a TSS scanning region for pol-II (5). The binding of general transcription factor proteins and histones to the TATA box facilitates the subsequent binding of pol-II which, along with several transcription factor proteins, constructs a transcription initiation complex that starts the mRNA synthesis from the TSS (13, 14). The nucleotide composition of different regions in the core promoter also has a strong influence on the sensitivity of the promoter. Studies have shown promoters with A/T- or T/C-rich PIC regions have higher sensitivities than promoters containing G/C-rich sequences (7). The position of the different regions of the core promoter is illustrated in figure 1.

**Figure 1:**
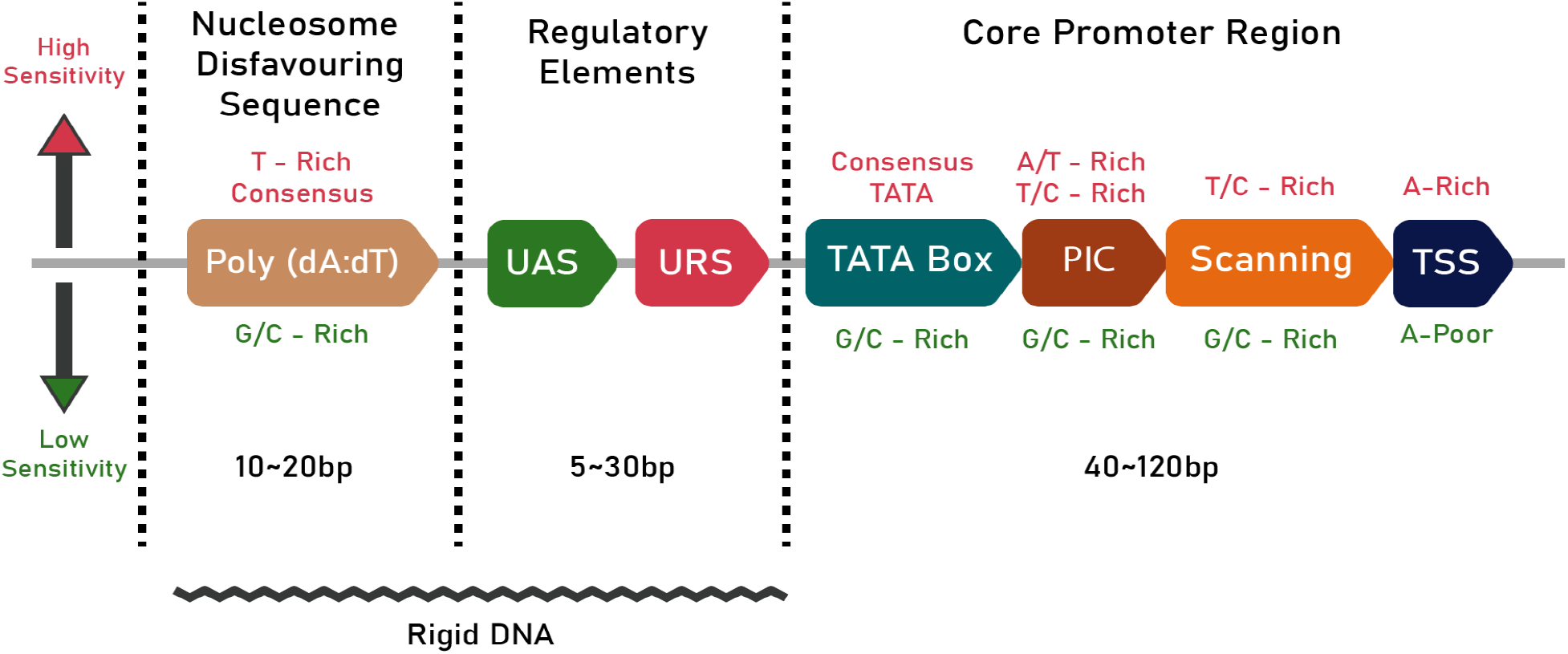
A schematic of the promoter architecture in S Cerevisiae. The text in Crimson (top) denotes the conditions necessary for high sensitivity while the green text (bottom) denotes the conditions for lower sensitivity. The length of the different regions of the promoter are also given in bp. The jagged line at the bottom denotes the part of the promoter that is rigid in nature.

The UAS and URS are the regulatory components of a promoter and are located upstream to the core promoter region. UASs and URSs act as binding sites for transcription activators and repressors, respectively. The UAS enhances gene expression, provides additional stability and plays a role in regulating the PIC formation process (8,9). The UASs and URSs in *S. cerevisiae* are typically 10 bp long but can vary from 5 to 30 bp in length (10).

The disfavouring nucleotide sequence is a stretch of DNA that decreases nucleosome occupancy to facilitate transcription (11). Poly(dA:dT) tract, a homopolymeric stretch of deoxyadenosine nucleotides, is a well-known nucleosome-disfavoring sequence commonly present in promoters (12).

The structural properties of the promoter are vital for successful transcription. The flexibility of the promoter should be optimal to make sure that the binding sites are accessible and properly positioned to enable their recognition by transcriptional machinery (13). In this regard, the bendability or the propensity of each trinucleotide to bend is essential (14). Existing studies indicate the presence of regions of low bendability about 100-200 bp upstream to the start codon (15) (illustrated in figure 1 by a jagged line). Studies also indicate that the low bendability is caused by a combination of A/T richness and di- and tri-nucleotide composition (16).

Promoters in *S. cerevisiae* can be either constitutive, that are relatively unaffected by internal and external signals and maintain stable levels of transcription, or inducible, which are capable of initiating a drastic change in transcriptional levels in response to specific stimuli. These stimuli, called inducers, range from molecules such as metabolites, amino acids and sugars, to metal ions and environmental factors like pH and stress (17,18). Using endogenous promoters of *S. cerevisiae* for synthetic biology applications has disadvantages owing to an insufficiency of well-characterized promoters (19,20). Thus, it is of utmost importance to characterize and quantify the strength of various *S. cerevisiae* promoters and create a database of the same.

Previous work has established a two-step approach for the quantitative prediction of the strength of promoters in *Escherichia coli (E. coli)*, a prokaryotic model organism (21,22). The linear relationship between the total promoter score and the promoter strength is well established in *E. coli* (22,23). In this study we have presented a similar simplified model of the promoter strength in *S. cerevisiae* based on the promoter sequence.

## Methods

The core promoter sequences of 5117 promoters in *S. cerevisiae* were retrieved using the Sequence Retrieval in Tool Eukaryotic Promoter Database (EPD) (24). The core promoter sequence consisted of −49 to 10 sequences with reference to the Transcription Start Site (TSS).

A Position Frequency Matrix (PFM) was generated from the motif of all 5117 promoter sequences. The PFM was then converted to Position Weight Matrix (PWM) or Position Specific Scoring Matrix (PSSM) using the functions from biopython (25). The motif landscape was visualized using weblogo (26). The resulting PSSM was then used to calculate the “promoter score” for all the promoters in the downstream analysis.

Taking inspiration from the well established linear relationship between the total promoter score and the promoter strength in *E. coli* (23), we modeled the promoter strength using a linear model with the promoter score as,

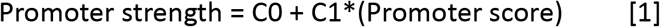

Lee *et al*. have characterized the strength of 19 constitutive promoters in *S. cerevisiae* using three fluorescence markers (Venus, mRuby2, and mTurquoise2) (27). We have used 18 of these promoter strengths in this study. The sequence for the promoter pREV1 was not found in EPD due to which we have dropped it from our analysis. The normalized fluorescence values folded over background were obtained from the authors of Lee *et al*. The log of these values gives the promoter strength. These were further divided by the strength of the strongest promoter (pTDH3) from Lee *et al*. We note that this step does not alter the linear relationship that is being tested but merely acts as a scaling. These values finally constitute the result space or the “Promoter strength” in equation [1].

We define a “segment score” which is simply the total score of a given segment of a given promoter as calculated from PSSM. This score is then divided by the highest score (corresponding to pTDH3) to obtain the feature space or “Promoter score” part of the equation [1].

We then performed Ordinary Linear Regression (OLS) using the statsmodels package in python. C0 and C1 were left as free parameters to obtain the best fit. The quality of fit was then assessed using reduced r-squared and F-statistic. The significance of model parameters were assessed using t-statistic. We also fitted a linear model to the residues to look for biases in the model. Scipy, Statmodel and Seaborn packages were used to perform, visualize, and test the linear regression (28).

## Results and Discussion

The 5117 native *S. cerevisiae* promoters from the EPD (29) that were included in this study represent a diverse population of transcriptional regulators. Motif analysis on this set revealed that the promoters were diverse in terms of nucleotide composition. We see that the conservation along the entire promoter length is low, except for the TSS. This is shown in figure 2.

**Figure 2:**
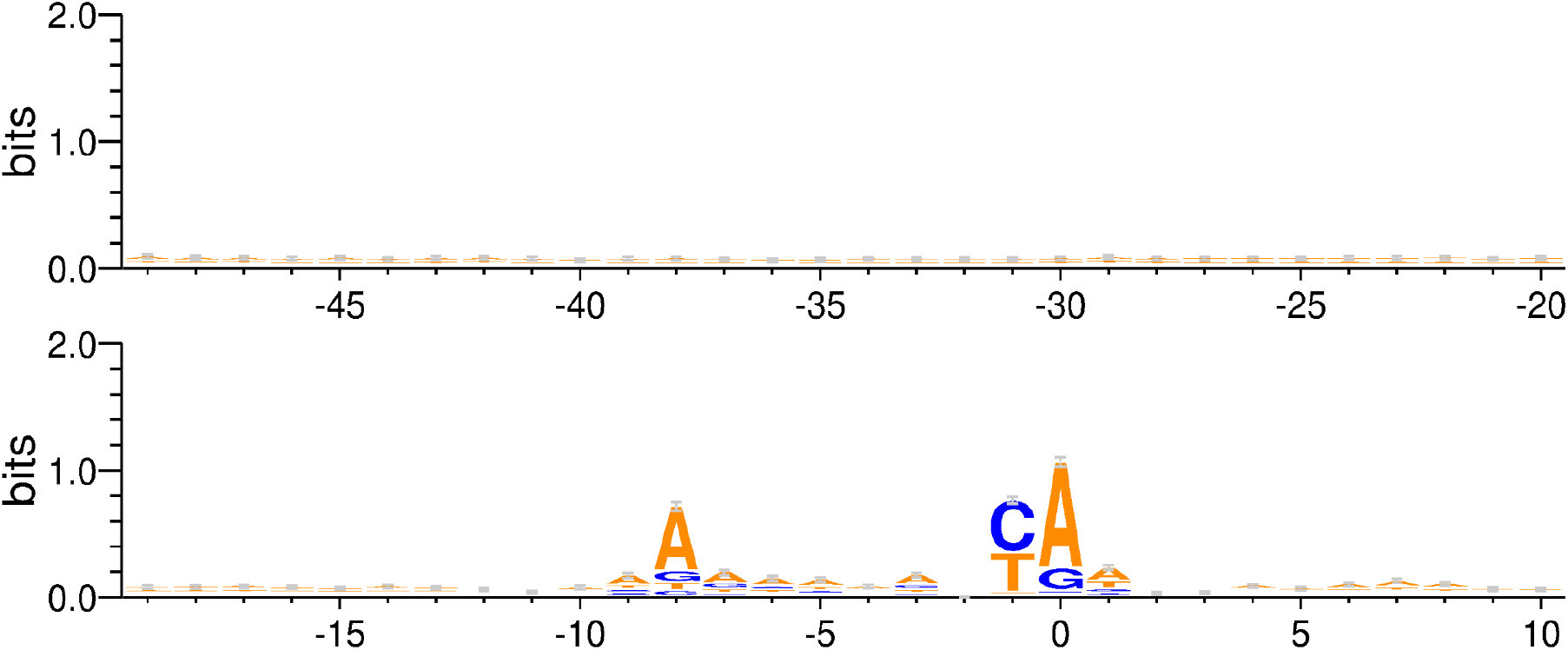
Motif logo generated from all 5117 Saccharomyces cerevisiae promoters from EPD.

There are two broad types of segments of the promoter that can be considered for modelling the linear relationship in equation [1]. These are based on the −49 to 10 motif shown in figure 2. We explored each possibility in detail while carefully considering their biological implications.

### -49 to X region

We sought to determine the shortest sequence that could satisfy the theoretical predictions and experimental results. One end of the segment, thus, was fixed at −49 and nucleotides were added stepwise towards the TSS until a nucleotide position ‘X’. Scores of the segments thus obtained were used to perform a linear regression described by the equation [1]. The quality of fit estimators for each such regression is shown in figure 3. It was observed that the quality of fit generally improves as more and more nucleotides are added. This is indicated by larger R-squared values and lower p-values for F-statistic. A saturation is reached at X=-1 after which adding more nucleotides does not improve the quality of fit by an appreciable amount as shown in figure 3 (and supplementary figure 1). This saturation is sustained until X=10, indicating that the quality of fit from −49 to −1 and −49 to 10 is mostly similar (Supplementary figures 4,5,6). Figure 4 shows the plot of normalized −49 to 10 scores and normalized mRuby2 fluorescence along with the best fit model. We see that the residues for this model are randomly distributed around 0, indicating that the errors are uncorrelated, and the quantile-quantile plot shows that errors are normally distributed. Similar trends were observed using Venus and mTurquoise2 fluorescence as seen in supplementary figures 2 and 3.

**Figure 3:**
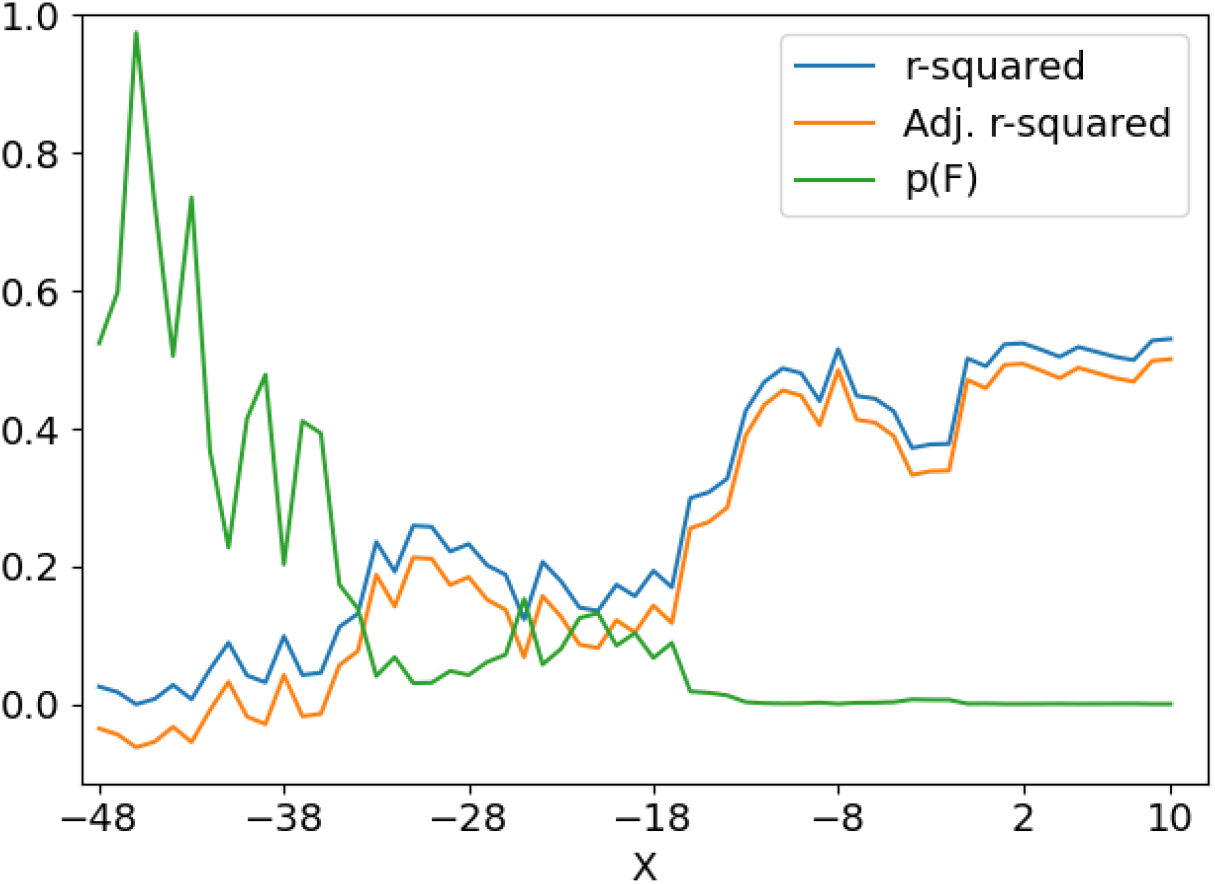
Various fit statistics for the linear regression of segment scores against the mRuby2 fluorescence. One of the ends of the promoter is fixed at −49 and nucleotides are added on the other end towards TSS. The values of R-squared, Adj. R-squared, and p-value for F-statistic are tabulated in supplementary table 1. Similar plots for Venus and mTurquoise2 fluorescence are given in supplementary figure 1.

**Figure 4:**
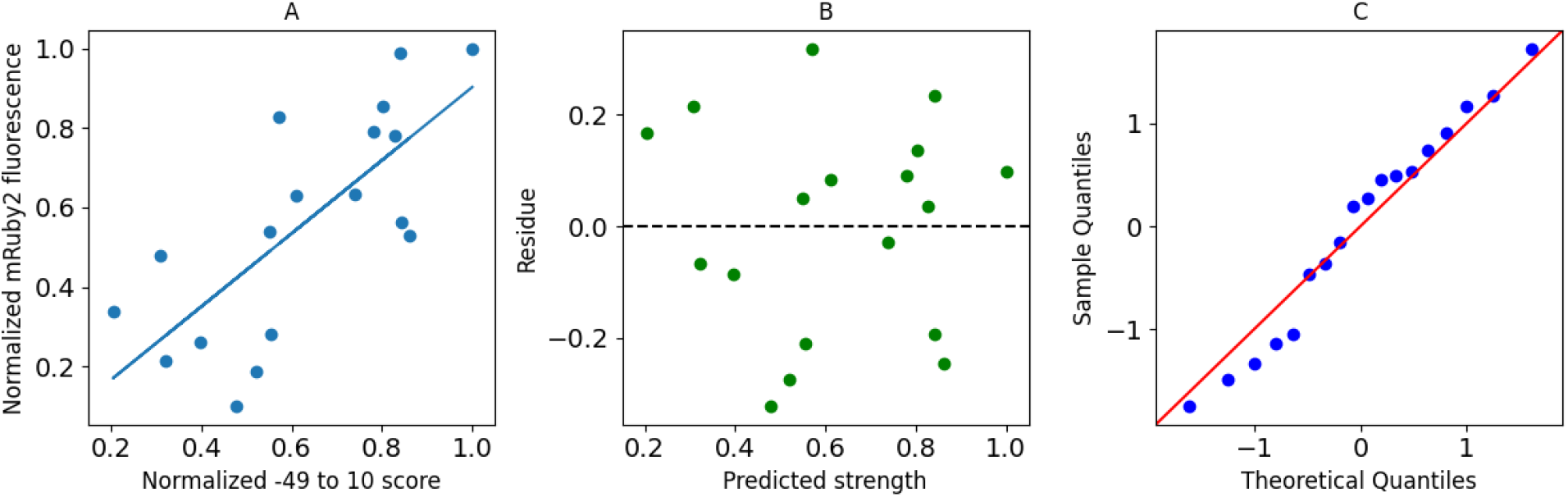
(A) Plot of normalized −49 to 10 score and normalized mRuby2 fluorescence along with the best fit model. (B) Residues obtained from the best fit model. (C) Quantile-Quantile plot of residues against normally distributed theoretical quantiles.

Since the −49 to 10 sequence was readily available in the EPD, further analysis focussed on this 60 bp fragment to ease integration with the EPD.

### -9 to 1 region

Motif analysis shown in figure 2 showed that the −9 to 1 region to be the most conserved stretch. Previous work by Berg & von Hippel suggests that conserved sequences contribute significantly to binding specificity of the promoter region (23). As mentioned earlier, a high binding specificity is indicative of high promoter strength. Consequently, we sought to determine whether this hypothesis holds true for the promoters included in our analysis. The quality of fit observed in this case, however, is extremely poor as seen in the figure 5. The p-value of F-statistic is 0.54 for regression using mRuby2 strengths. Therefore, we can not reject the null hypothesis that there is no relationship between −9 to 1 score and promoter strength. This demonstrates that the above argument is not a dominant mechanism in defining the promoter strength.

**Figure 5:**
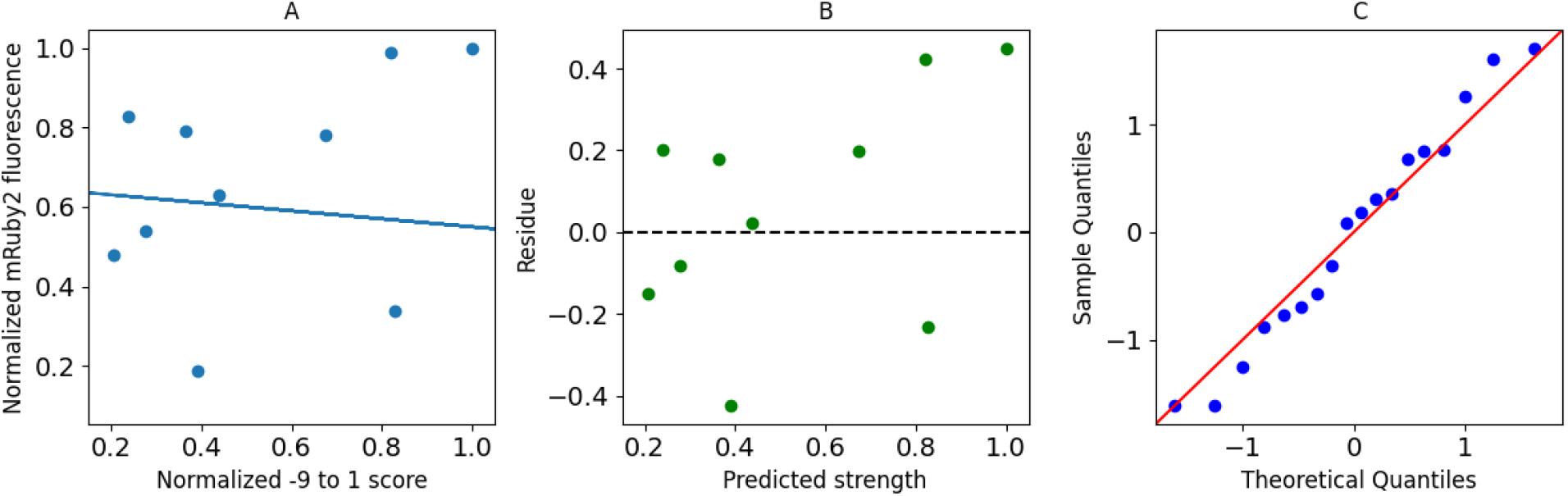
(A) Plot of normalized −9 to 1 score and normalized mRuby2 fluorescence along with the best fit model. (B) Residues obtained from the best fit model. (C) Quantile-Quantile plot of residues against normally distributed quantiles.

We found that the value of intercept C0 in the model is close to zero for the best fit model and the value of slope C1 was between 0.8 and 1.1. Figure 6 shows the best fit values of C0 and C1 for different fluorescence along with their errors. Values of C0 and C1 obtained using mTorquoise2 fluorescence were slightly different from those obtained using Venus or mRuby2 fluorescence. These differences likely stem from the stochastic gene expression and noisy fluorescence signals in the experiments. However, the error bars on these parameters do overlap. The mean values of C0 and C1 are −0.07 and 0.93 respectively.

**Figure 6:**
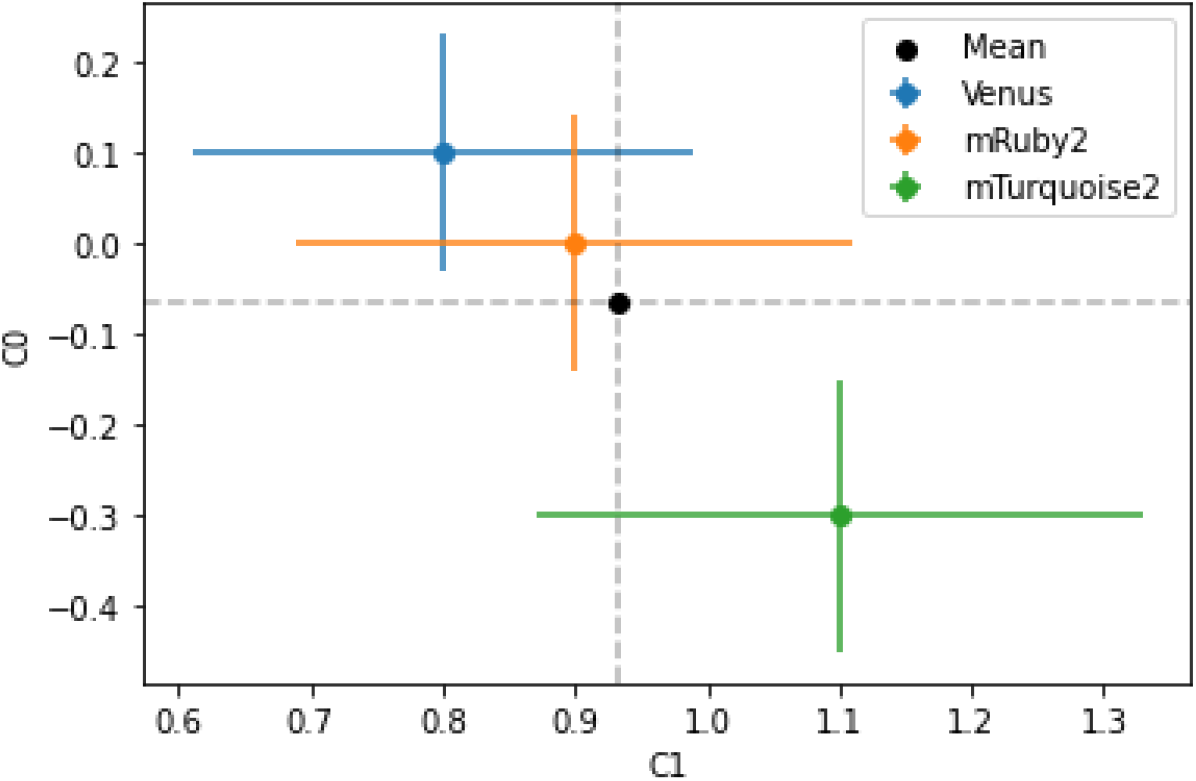
The best fit values of C0 and C1 obtained using different fluorescence data as a proxy for promoter strength. Black dot shows the mean value of C1 and C2 weighted by error bars. The data corresponding to this plot can be found in supplementary table 2.

**Figure 7:**
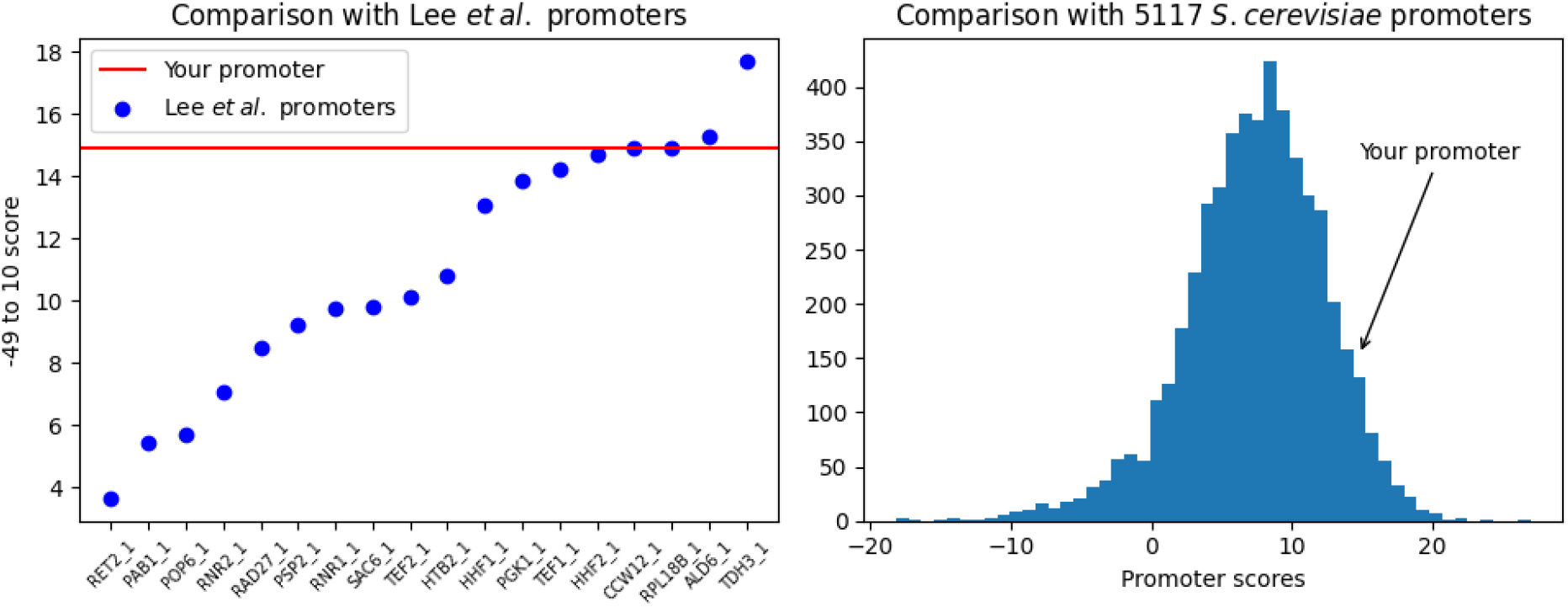
Example figures from the app. (a) −49 to 10 score of the user’s promoter is shown as a horizontal line on a plot of scores of the characterized promoters from Lee et al. (b) Arrow shows the score of the user’s promoter in reference to scores of all 5117 S. cerevisiae promoters in EPD.

We have developed an open source free to use standalone tool and a website to use our findings to predict the strength of *S. cerevisiae* promoters. The standalone tool can be found at https://github.com/DevangLiya/QPromoters and the website can be found at https://qpromoters.com/. Users can either enter the −49 to 10 sequence of the promoter of interest or they can retrieve this sequence directly from EPD by entering the EPDnew ID of the promoter. The tool then outputs the promoter score, promoter score normalized by pTDH3 score, promoter strength using the model described by equation [1], a plot showing the location of the user’s promoter with reference to the 18 characterized promoters, and a histogram showing the location of the user’s promoter with reference to all 5117 EPD promoters (29). An example of the figure returned by the program is shown in figure 5.

## Conclusion

Our study attempted to define the minimal promoter region representative of the strength of the entire promoter in *S. cerevisiae*. To this end, we analysed the core promoter region of −49 to 10, with respect to TSS, to find a simple correlation between the score of this region and the experimental promoter activity. After analysing the core promoter region, it was evident that there is a correlation between the promoter score and experimental promoter strength. Particularly, the −49 to 10 region of the core promoter was seen to be the best predictor of the promoter strength. We also observed a similar quality of fit between the −49 to −1 and −40 to 10 regions. The biological basis and significance of this sustained quality of fit needs further investigation. In addition to these findings, we have developed an open source, free-to-use tool to predict the promoter strength of unknown promoters in *S. cerevisiae*.

Using computational tools to determine essentiality of genes and strength of gene regulatory elements is of significant use in synthetic biology as it allows for back-tracking in case of failure and is less time-consuming (30). Our work is also relevant in the efforts that are being made to construct the minimal yeast genome. Construction of such a genome would be an ideal proof-of-concept to facilitate the precise engineering of genomes tailored to meet particular environmental or functional requirements. Consequently, these in-silico methods can precede and lower the risk of failure in in vivo experiments. Moreover, our web service could be useful in characterizing the strength of existing promoters in the EPD as well as predicting the strength of other engineered *S. cerevisiae* promoters on the basis of promoter sequence.

## Conflict of interest

The authors declare no conflict of interest.

## Funding

The authors did not receive funding from any source for this work and was carried out of scientific interest.

## Acknowledgement

The authors would like to thank Shubham Kumar Sinha and Swaroopa Nakkeeran from Indian Institute of Science Education and Research, Mohali for their comments and suggestions throughout the work.

## Authors contribution

AKJ conceptualized and designed the project. DHL curated the data and worked on the formal analysis to quantify the promoter strength. NMA and NB did the promoter landscape analysis and visualizations. MP designed web implementation and provided technical assistance. AKJ, SS, DHL, ME wrote the manuscript. All the authors have read and approved this manuscript.

## Supplementary Figures

**Supplementary Figure 1:**
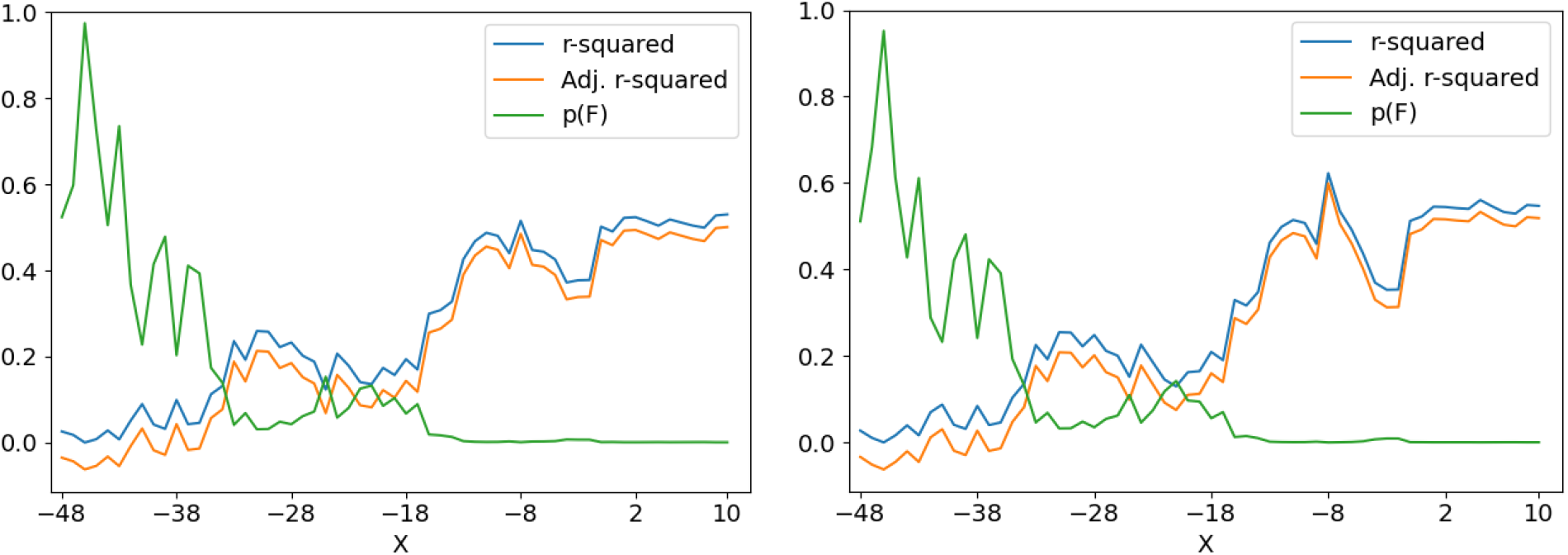
Various fit statistics for the linear regression of segment scores against the (A) Venus fluorescence and (B) against the mTurquoise2 fluorescence. One of the ends of the promoter is fixed at −49 and nucleotides are added on the other end towards TSS. The values of R-squared, Adj. R-squared, and p-value are tabulated in supplementary table 1.

**Supplementary Figure 2:**
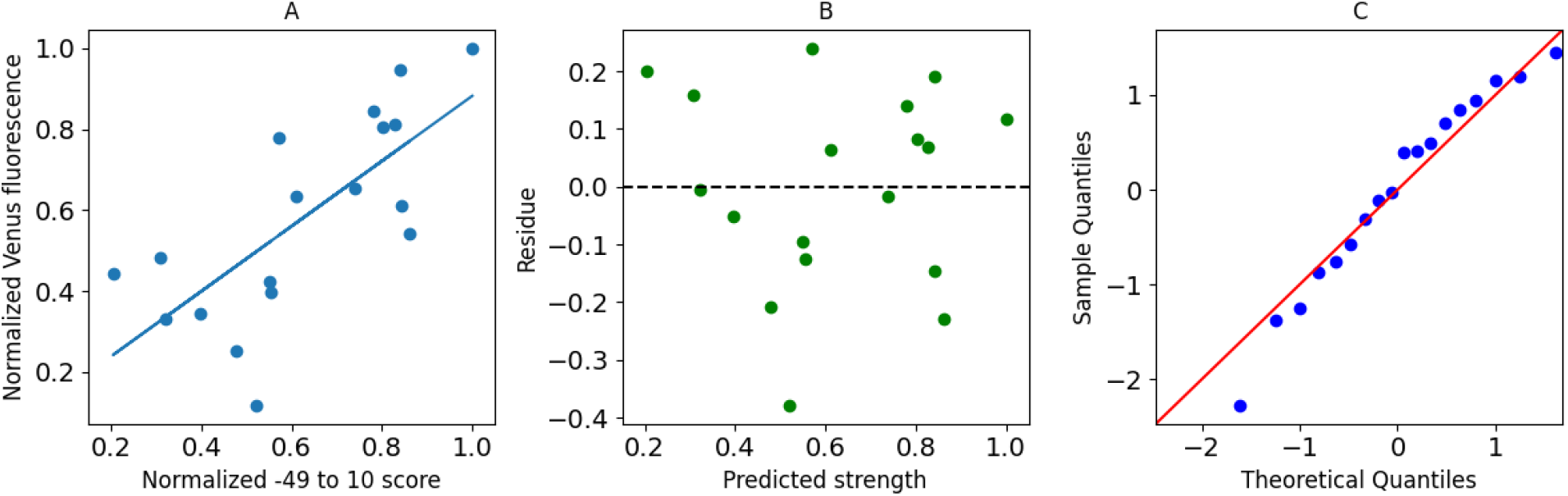
(A) Normalized promoter score is plotted against normalized Venus fluorescence. Blue line shows the best fit model. (B) Residues from the model. (C) Quantile-Quantile plot for normally distributed quantiles.

**Supplementary Figure 3:**
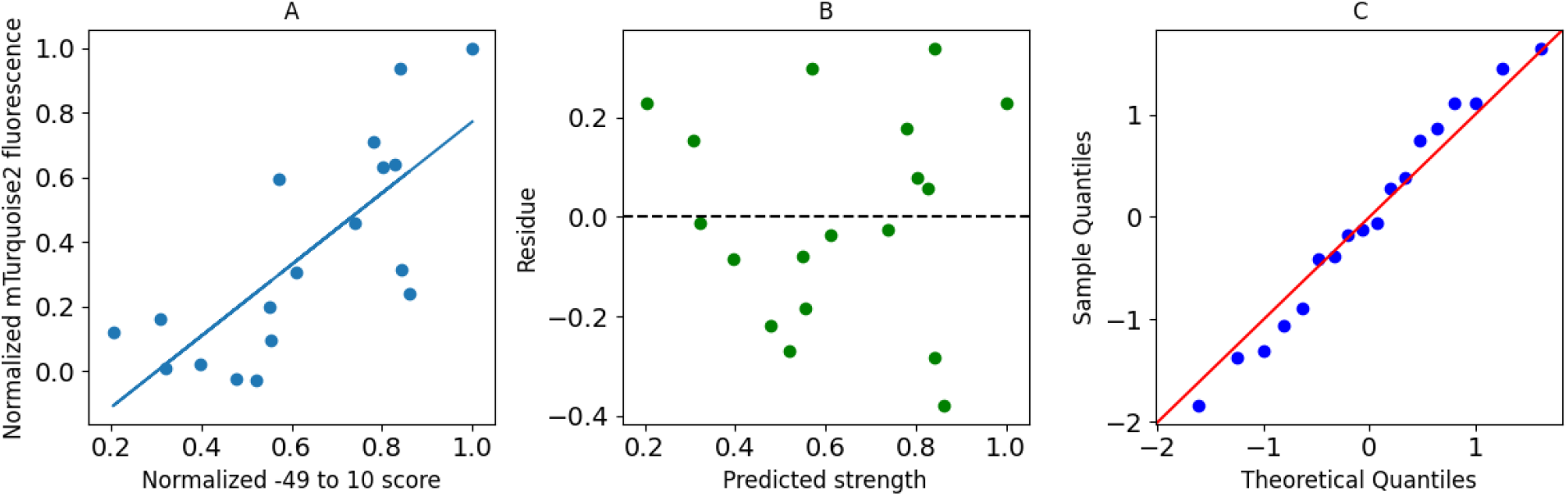
(A) Normalized promoter score is plotted against normalized mTurquoise2 fluorescence. Blue line shows the best fit model. (B) Residues from the model. (C) Quantile-Quantile plot for normally distributed quantiles.

**Supplementary Figure 4:**
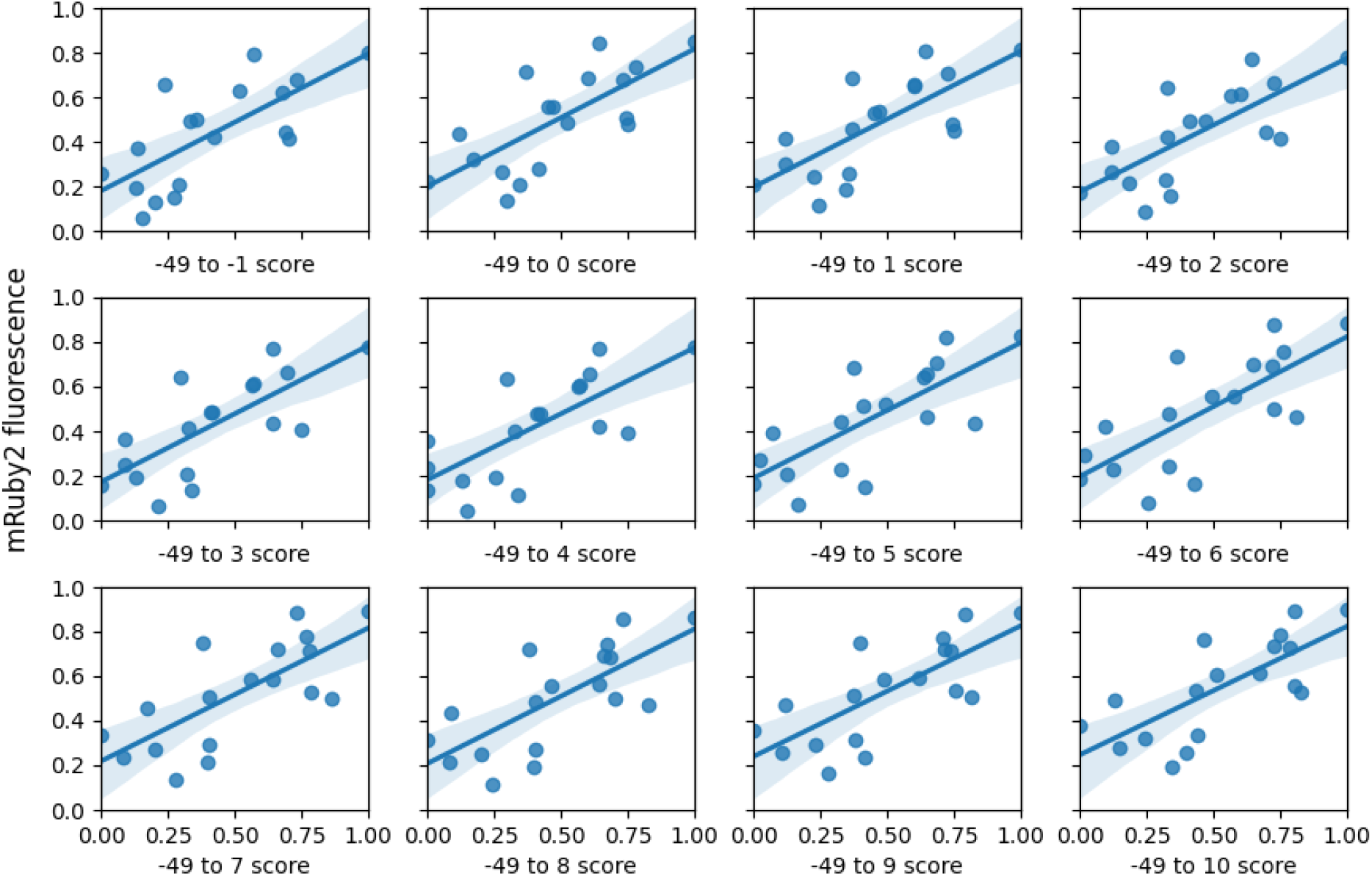
Normalized mRuby2 fluorescence plotted against the normalized promoter score. The solid blue line shows the best fit line along with 95% interval. The promoter scores of different panels are calculated using only −49 to X region (where X varies from −1 to 10) with respect to TSS. See section 1 in results and discussion for more information.

**Supplementary Figure 5:**
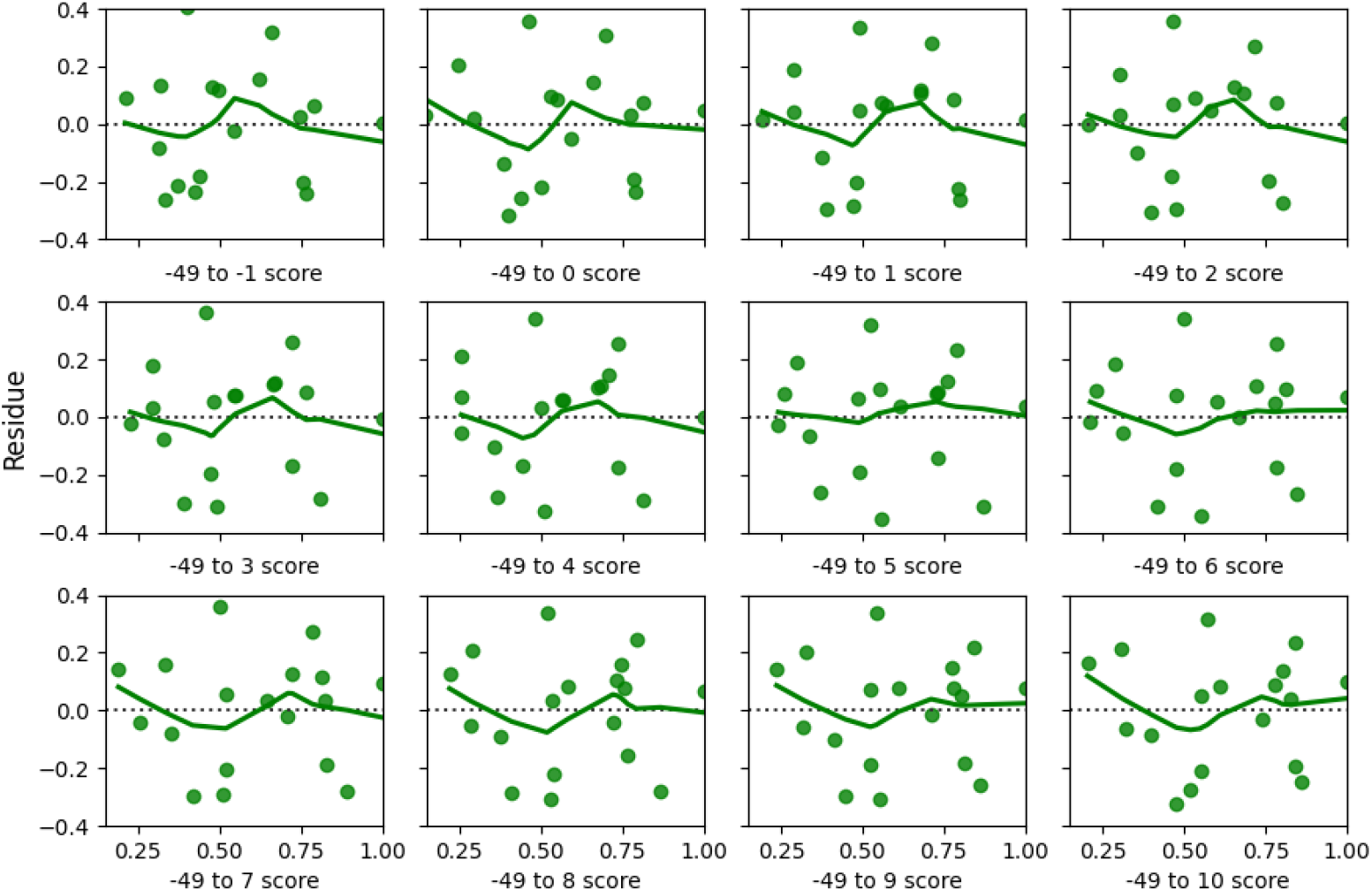
This figure shows the residual plots corresponding to the fits in supplementary figure 4.

**Supplementary Figure 6:**
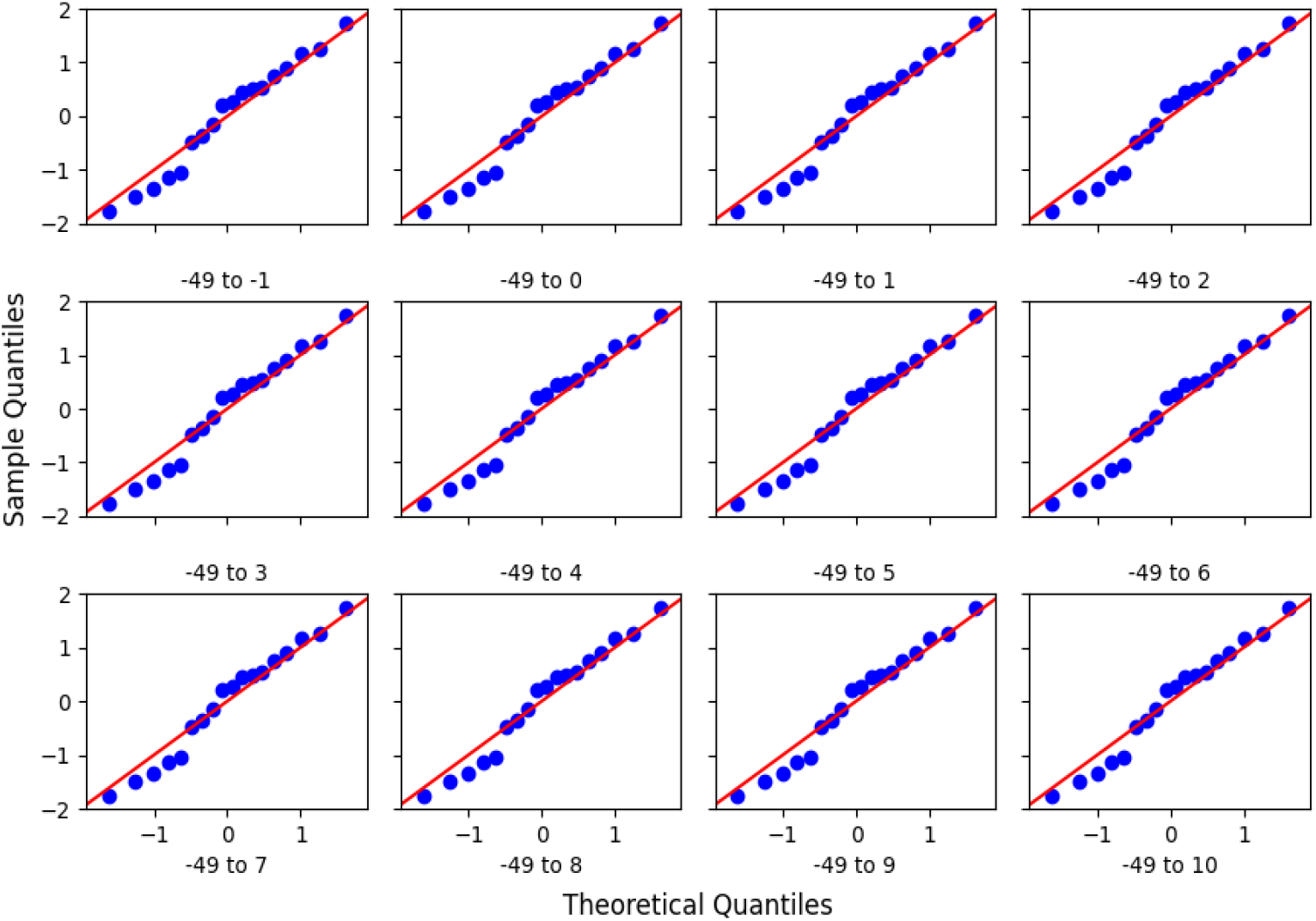
This figure shows the Quantile-Quantile plots for residues corresponding to the fits in supplementary figure 4 plotted against normally distributed theoretical quantiles.

## Supplementary Tables

**Supplementary table 1:**
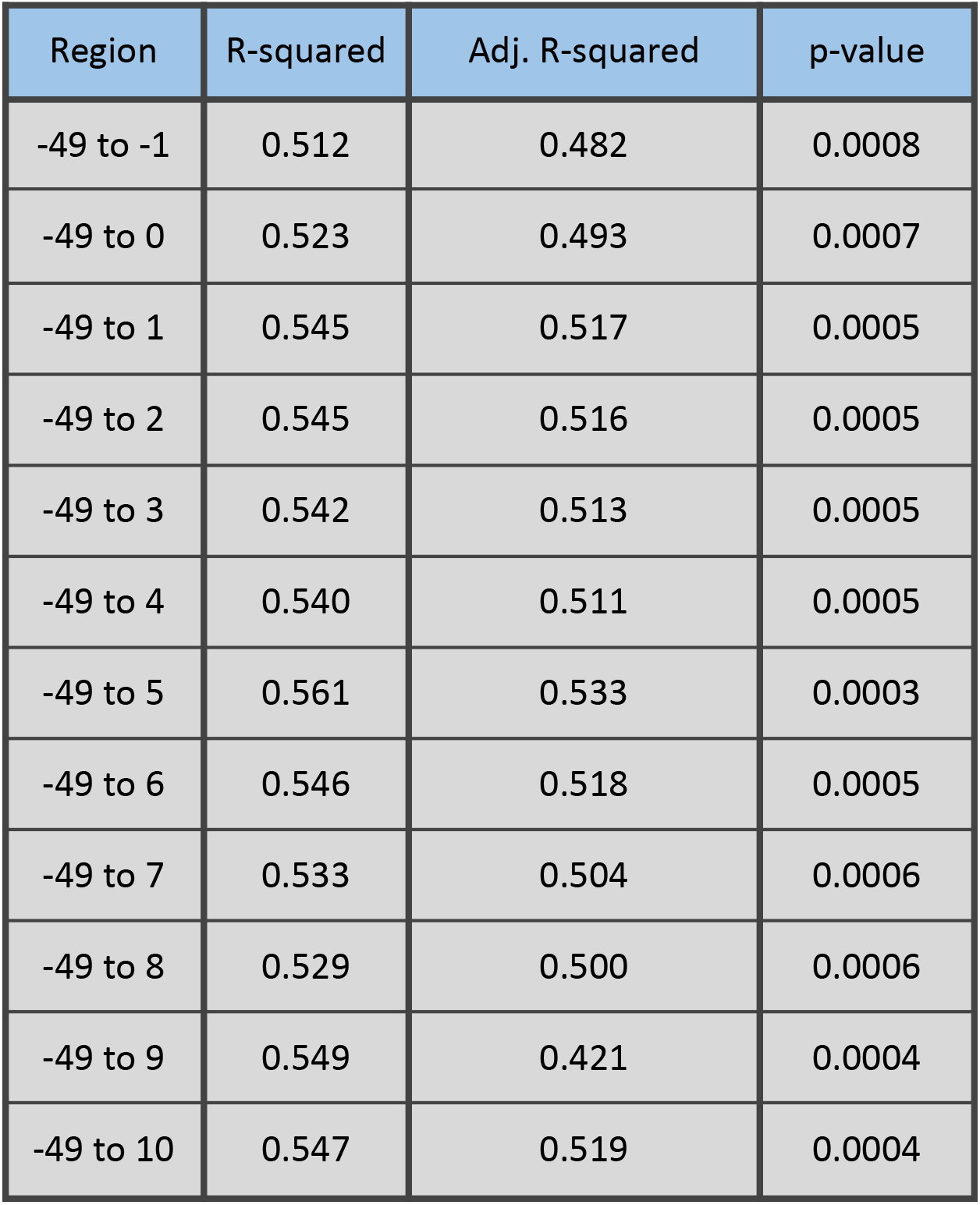
Assessment of the quality of fit using mRuby2 fluorescence values and promoter score when one of the ends of the promoter is fixed at −49 and nucleotides are added on the other end.

**Supplementary Table 2:**
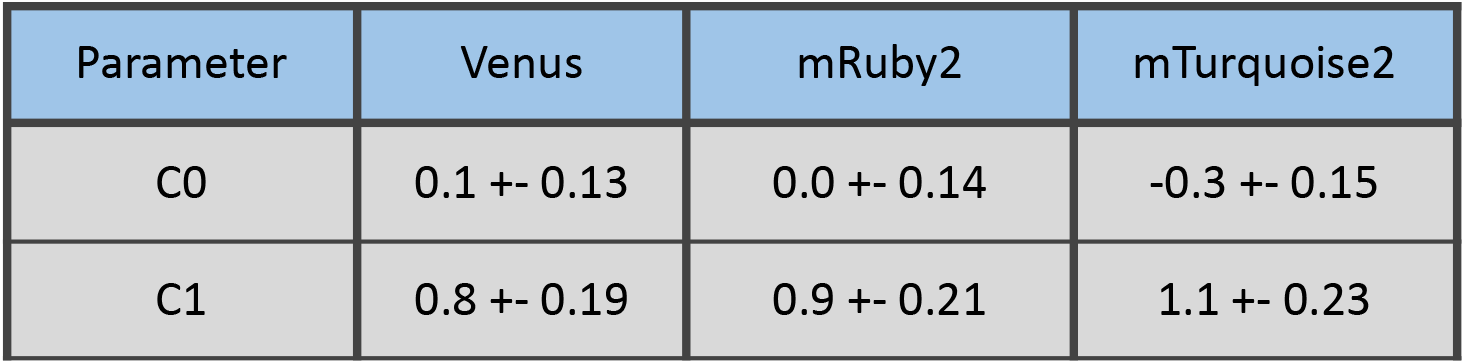
Best fit values of model parameters using different fluorescence values as indicator for experimental promoter strength.

## Notes

### Competing Interest Statement

The authors have declared no competing interest.

