## Supplementary figures and tables for "QPromoters: Sequence based prediction of promoter strength in *Saccharomyces cerevisiae*"

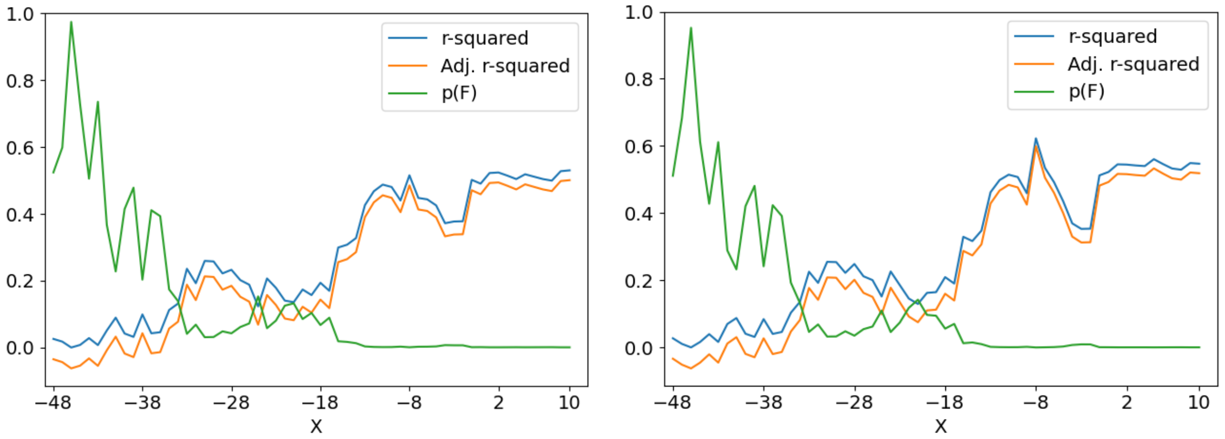


***Supplementary Figure 1:*** *Various fit statistics for the linear regression of segment scores against the (A) Venus fluorescence and (B) against the mTurquoise2 fluorescence. One of the ends of the promoter is fixed at -49 and nucleotides are added on the other end towards TSS. The values of R-squared, Adj. R-squared, and p-value are tabulated in supplementary table 1.*

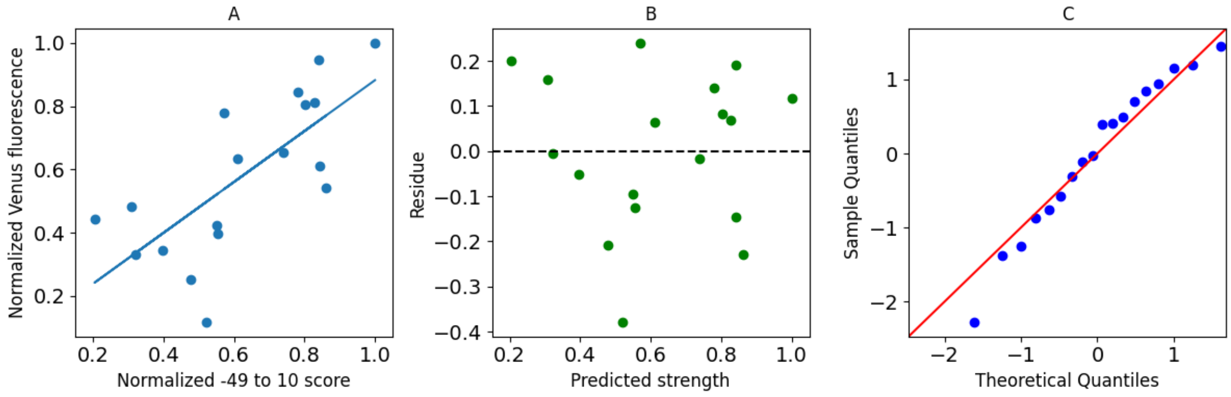


***Supplementary Figure 2:*** *(A) Normalized promoter score is plotted against normalized Venus fluorescence. Blue line shows the best fit model. (B) Residues from the model. (C) Quantile-Quantile plot for normally distributed quantiles.*

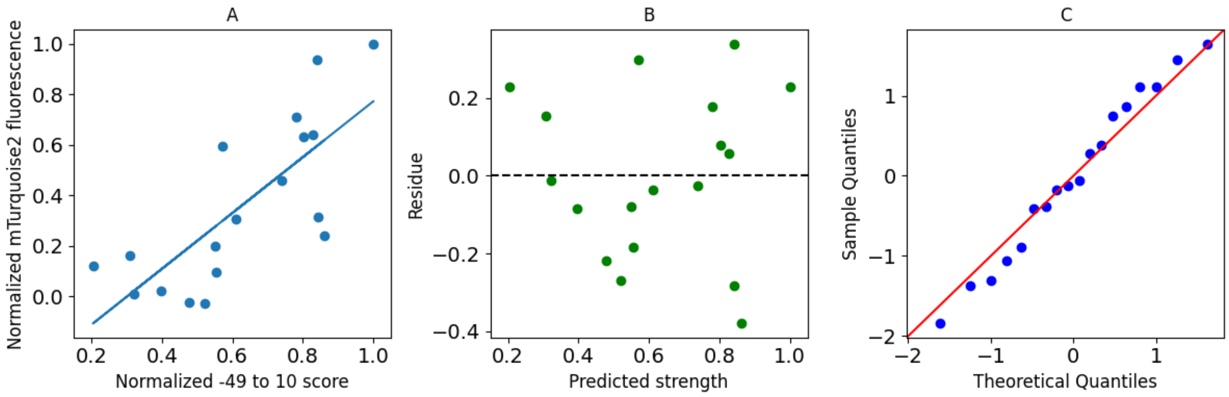


***Supplementary Figure 3:*** *(A) Normalized promoter score is plotted against normalized mTurquoise2 fluorescence. Blue line shows the best fit model. (B) Residues from the model. (C) Quantile-Quantile plot for normally distributed quantiles.*

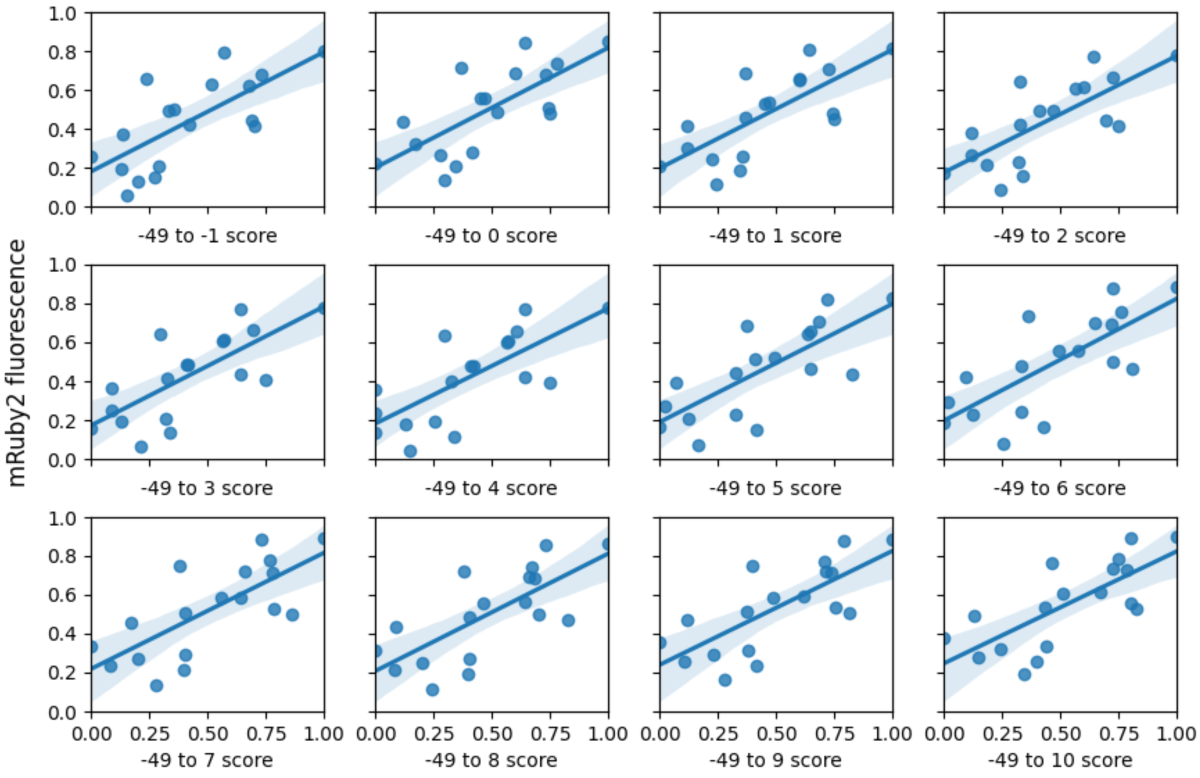


***Supplementary Figure 4****: Normalized mRuby2 fluorescence plotted against the normalized promoter score. The solid blue line shows the best fit line along with 95% interval. The promoter scores of different panels are calculated using only -49 to X region (where X varies from -1 to 10) with respect to TSS. See section 1 in results and discussion for more information.*

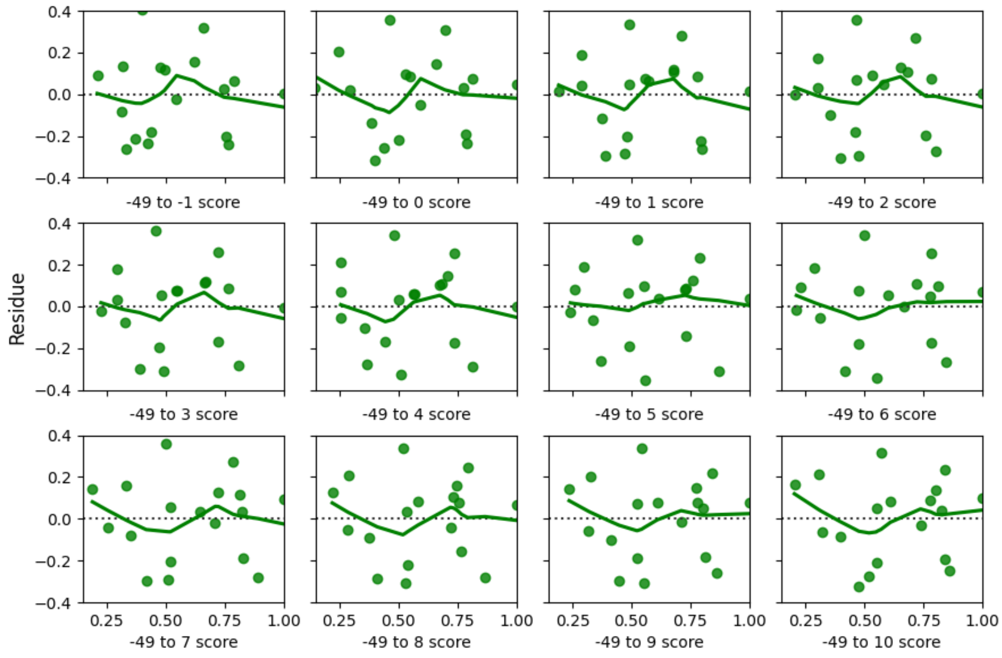


***Supplementary Figure 5:*** *This figure shows the residual plots corresponding to the fits in supplementary figure 4.*

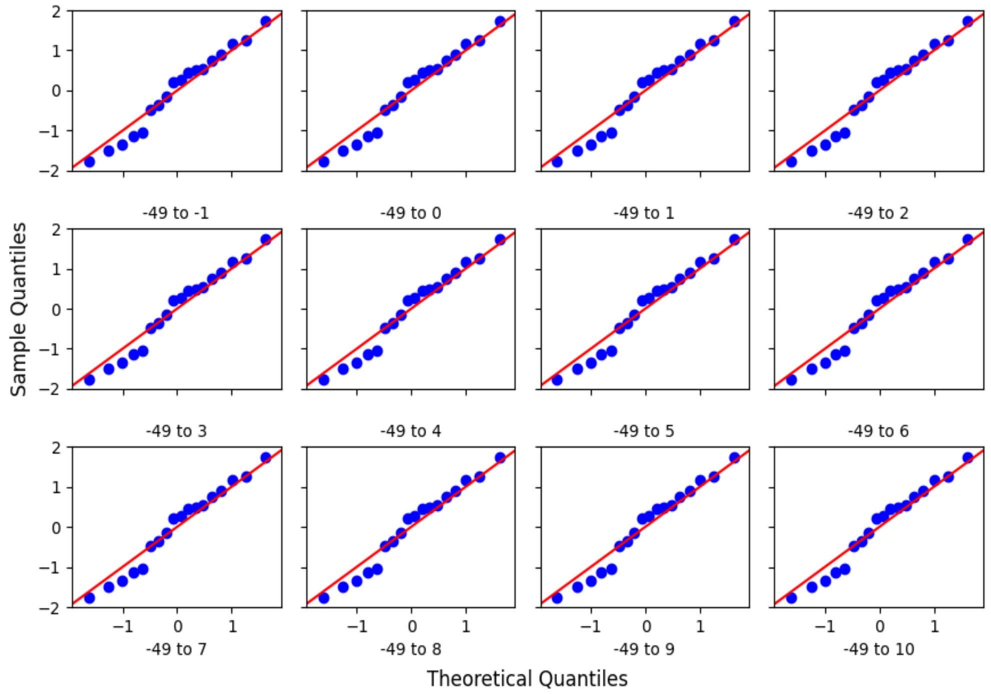


***Supplementary Figure 6:*** *This figure shows the Quantile-Quantile plots for residues corresponding to the fits in supplementary figure 4 plotted against normally distributed theoretical quantiles.*

### Supplementary Tables

| Region | R-squared | Adj. R-squared | p-value |
|---|---|---|---|
| -49 to -1 | 0.512 | 0.482 | 0.0008 |
| -49 to 0 | 0.523 | 0.493 | 0.0007 |
| -49 to 1 | 0.545 | 0.517 | 0.0005 |
| -49 to 2 | 0.545 | 0.516 | 0.0005 |
| -49 to 3 | 0.542 | 0.513 | 0.0005 |
| -49 to 4 | 0.540 | 0.511 | 0.0005 |
| -49 to 5 | 0.561 | 0.533 | 0.0003 |
| -49 to 6 | 0.546 | 0.518 | 0.0005 |
| -49 to 7 | 0.533 | 0.504 | 0.0006 |
| -49 to 8 | 0.529 | 0.500 | 0.0006 |
| -49 to 9 | 0.549 | 0.421 | 0.0004 |
| -49 to 10 | 0.547 | 0.519 | 0.0004 |

***Supplementary table 1:*** *Assessment of the quality of fit using mRuby2 fluorescence values and promoter score when one of the ends of the promoter is fixed at -49 and nucleotides are added on the other end.*

| Parameter | Venus | mRuby2 | mTurquoise2 |
|---|---|---|---|
| C0 | 0.1 +- 0.13 | 0.0 +- 0.14 | -0.3 +- 0.15 |
| C1 | 0.8 +- 0.19 | 0.9 +- 0.21 | 1.1 +- 0.23 |

***Supplementary Table 2:*** *Best fit values of model parameters using different fluorescence values as indicator for experimental promoter strength.*
